# Agricultural intensification alters bee community composition and functional diversity across tropical heterogeneous landscapes

**DOI:** 10.64898/2026.11.26.769391

**Authors:** Dona roy, Indranil Samajpati, Aditi dutta, Parthiba Basu, Rita Kundu

**Affiliations:** Ecology Research Unit, Department of Zoology, University of Calcutta, 35, B.C. Road, Kolkata 700019, India; Advanced Cell Biology Laboratory, Department of Botany, University of Calcutta, 35, Ballygunge Circular Road, Kolkata 700019, West Bengal, India

**Keywords:** Pollination, Agricultural intensification, Landscape, Bee assemblage, Network

## Abstract

Agricultural intensification and landscape features influence pollinator diversity globally; however, their impact on site-pollinator interactions and the functional diversity of bees is scarce in the literature, especially in tropical agricultural systems. To meet the knowledge gap, we revealed how pesticide intensity, crop diversity, and semi-natural vegetation influence bee communities across high and low agricultural intensification systems across 19 sites in the Terai region of West Bengal, India. Pesticide intensity negatively influenced bee abundance and functional richness, indicating environmental filtering. Site-species interaction network revealed that the network structure was more nested in high AI sites, while less specialized, suggesting these sites are dominated by disturbance-tolerant generalist species. Whereas, Low AI sites showed more heterogeneous and specialized interactions. Contrastingly, in both high and low AI sites, *Lassioglossum* sp. 2 species showed the highest normalized degree. Bee diversity and functional dispersion of bees within crops field negatively associated with natural vegetation, indicating that surrounding natural vegetation might act as an alternative foraging refuge. Our findings show that agricultural intensification alters bee community structure, reduces ecological complexity, and may compromise long-term pollination stability despite the persistence of generalist pollinators.

## 1. Introduction

Agricultural intensification (AI) in recent decades has led to widespread declines in global farmland biodiversity. This includes a rapid decline in wild bee populations, which are vital for pollinating both crops and wild plants (Robinson and Sutherland, 2002; Klein et al., 2007; Potts et al., 2010; Kennedy et al., 2013). Drivers of AI, including habitat loss and fragmentation, reduced landscape heterogeneity, monoculture expansion, and increased pesticide use (Steffan-Dewenter and Tscharntke, 2000; Fahrig et al., 2003; Geiger et al., 2010) individually, as well as interactively affects local bee populations. Such combined effects are particularly relevant in tropical agricultural systems, where smallholder farming, heterogeneous land use, and unregulated management practices create complex environmental gradients (Tscharntke et al., 2012). Evaluation of multiple components of agricultural intensification simultaneously therefore, provides a more comprehensive understanding of how bee communities respond to landscape modification and environmental change. Due to the essential role of wild bees in maintaining crop production and ecosystem functioning, understanding how agricultural intensification affects bee communities has become a major concern for both conservation efforts and sustainable agricultural practices (Klein et al., 2007; Potts et al., 2010).

Apart from AI, landscape features also influences bee communities in agricultural farms. Landscape-level factors such as Semi-natural habitats (SNH), crop diversity, and fallow lands provide nesting resources and continuous floral availability that support diverse bee assemblages (Kennedy et al. 2013; Garibaldi et al. 2011; Raderschall et al., 2020; Lasway et al., 2022). These factors can counteract the detrimental effects of AI, ultimately increasing resilience of local pollinator assemblages (Brittain and Potts, 2011; Nicholls & Altieri, 2012). Landscape heterogeneity, particularly in tropical agricultural systems characterized by smallholder farms and mixed cropping systems, may buffer the negative effects of intensification by maintaining resource diversity (Priyadarshana et al., 2024). However, the responses of bee communities to multiple components of agricultural intensification and landscape features, including pesticide input, SNH, and crop diversity, remain insufficiently understood, especially in tropical agroecosystems (Regmi et al., 2025; Chakraborty & Basu, 2025).

Moreover, simple community metrics such as species richness and abundance is often inadequate to understand community responses to environmental and anthropogenic changes. In such cases, functional trait approach provides more holistic insights into bee community responses (Coutinho et al., 2021). Functional traits such as body size, nesting behaviour, and foraging specialization determine how bee species respond to landscape modifications and influence ecosystem functioning (Coutinho et al., 2021; Felderhoff et al., 2023). Agricultural intensification can filter bee communities by favouring disturbance-tolerant species, thereby reducing functional diversity and altering ecosystem resilience (Chakraborty & Basu, 2025). Therefore, evaluating functional diversity indices, including functional richness, evenness, and dispersion, can improve understanding of how intensification affects ecosystem stability and pollination services (Chakraborty & Basu, 2025; Dutta et al., 2024).

Beyond functional diversity, ecological network approaches provide additional insights into bee community structure and the associations between species and their ecological environment (Basu et al., 2016; Marini et al., 2019; Dutta et al., 2024). Network analysis allows community patterns to be examined through the relationships among species and the habitats they occupy, providing information on community organization and potential responses to environmental changes (Lami et al., 2021; Marini et al., 2019). In this study, we used species-site networks to examine how bee species are associated with different sites across agricultural landscapes and how these associations vary along an agricultural intensification gradient. Network structure was characterized using topological metrics, including nestedness, modularity and specialization, which provide insights into species distribution, habitat associations and the organization of bee communities.

Theoretical studies suggest that bee-site networks in landscapes with low agricultural intensification may exhibit lower nestedness and greater specialization, reflecting stronger associations between bee species and particular sites (Fletcher et al., 2013). In contrast, increasing agricultural intensification and landscape simplification may favour widespread, disturbance-tolerant generalists over habitat-restricted species, potentially resulting in more nested network structures (Stouffer & Bascompte, 2011; Gámez-Virués et al., 2015). However, empirical studies simultaneously examining functional diversity and network structural changes across agricultural intensification gradients, particularly in tropical agricultural landscapes, remain limited (Dutta et al., 2024).

India’s agricultural landscapes, characterized by smallholder farms, mixed cropping systems, and high landscape heterogeneity, provide an ideal setting to evaluate the effects of agricultural intensification on bee communities (Basu et al., 2016). However, studies examining functional diversity and interaction networks in Indian agroecosystems remain limited. Moreover, these tropical agricultural landscapes are often influenced by multiple simultaneous stressors, making it essential to assess their combined impacts on bee communities and ecosystem functioning (Belsky and Joshi, 2019). Previous studies in Indian agroecosystems have shown that loss of SNH and increased AI alters bee community structure, functional diversity, and interaction networks in tropical systems (Dutta et al., 2024; Basu et al., 2016; Chakraborty & Basu, 2025). However, the findings are highly context dependant, with only a small number of studies spanning relatively few landscapes, despite India being one of the world’s most agroecologically diverse countries (Singh et al., 2021). Hence, more studies are essential for understanding these patterns for developing sustainable agricultural management practices that support pollinator conservation, especially in the different agroecological landscapes of India. In this study, we investigated how agricultural intensification influences bee community structure, functional diversity, and community structure in a heterogeneous agricultural landscape located in Northern Terai region of India. We examined the effects of pesticide input, semi-natural vegetation, and crop diversity on bee diversity and functional traits. We also observed how bee-site networks differ between high and low AI sites.

We hypothesized that:

- Bee abundance, diversity and functional diversity would decline with increasing agricultural intensification and pesticide input.
- Semi-natural vegetation and crop diversity would positively influence bee abundance, diversity and functional diversity.
- Landscapes with greater AI would exhibit greater nestedness but lower specialization in bee-site networks.

## 2. Methodology

### 2.1. Study site

This study was conducted in the district of Alipurduar of West Bengal, India. Alipurduar district is located in the northeastern corner of the state, extending from approximately 26°16′N to 26°52′N latitude and 89°00′E to 89°54′E longitude. The state is a part of the Eastern Dooars region, bordered by Bhutan, Assam, Cooch Behar, and Jalpaiguri (Saha and Das, 2024). The district has extensive but fragmented forest landscapes, such as Buxa Tiger Reserve, Rajabhatkhawa, Chilapata, and Raidak forests. The climate is tropical, with an average annual rainfall of approximately 3100 mm and temperature variations ranging from below 10 °C in winter to above 35 °C in summer (Ghosh et al., 2025). The main kharif crop in the region is rice. Winter crop (rabi) such as mustard is predominant in the region, while vegetables such as brinjal, tomato, cauliflower, cabbage, beans, cucumber, pumpkin, and chilli are also widely cultivated (Basak et al., 2023). The forests of Alipurduar are highly diverse and ecologically significant, encompassing dry deciduous, moist deciduous, semi-evergreen, and evergreen forest types. These forests are predominantly dominated by *Shorea robusta* (sal), along with other important tree species such as teak, sissoo, schima, simul, and a variety of evergreen taxa (Das, 2020; Barman et al., 2022). Despite this diversity, the landscape has become increasingly fragmented, with forests interspersed by tea plantations and human settlements. Since 1947, more than 20% of the original forest cover has been lost due to the expansion of plantations and other anthropogenic activities (Das, 2020; Barman et al., 2022). We selected 19 study sites (50m x 50m) in and around pollinator-dependent vegetable crop fields along an orthogonal gradient of semi-natural habitat and pesticide use intensity (Figure 1). Sites were spaced at least 1km from each other to maintain spatial independence.

**Figure 1.**
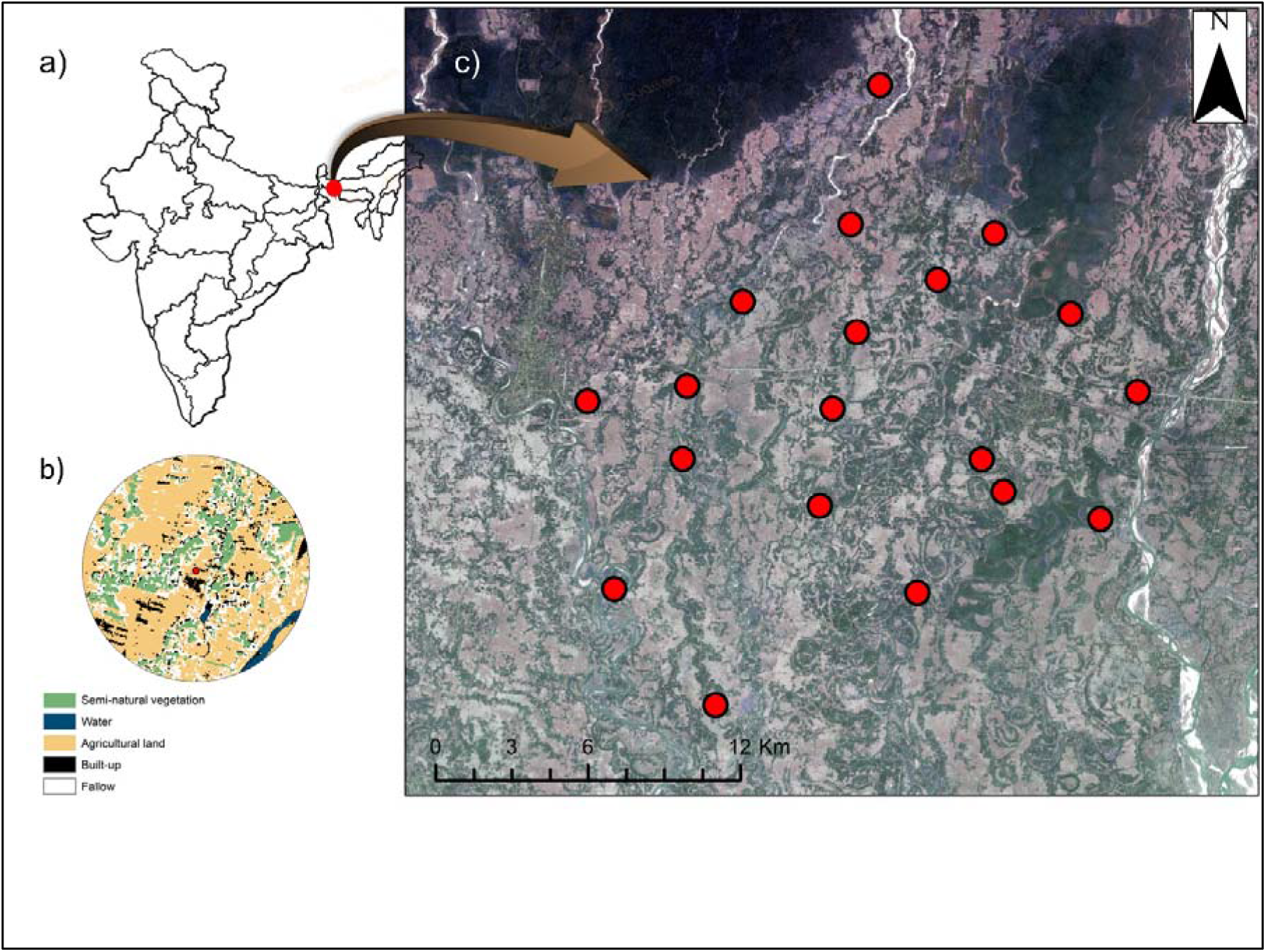
(a) Map of India with the study site highlighted in red. (b) Land Use Land Cover map of one of the study sites with a 500-m radius around focal fields. Semi-natural habitats are shown in green. (c) Satellite image of the study area with red dots indicating the locations of the study sites where the research was conducted.

### 2.2. Bee collection and identification

Bee sampling was conducted from November 2022 to March 2023 across the 19 study sites. Each site was sampled twice during the study period, with one complete sampling round encompassing all 19 sites in each sampling bout (total = 38 sampling days).The two sampling bouts were performed with a minimum gap of 15 days. Bee communities were quantified using a combination of passive (pan traps) and active (sweep netting) sampling methods.

In each sampling round, five pan trap units were installed randomly within agricultural farms, primarily consisting of seasonal vegetable crops such as eggplant, chilli, tomato, flat bean, and mustard, for passive bee sampling. At each site, a cluster of five trap units was established within a randomly selected 50 m × 50 m plot, with four trap units positioned at the corners of the plot and one placed at the centre (Figure 2). Coordinates of the pan trap stations were taken using GPS (Garmin, e-treks-30). We placed the pantraps of three different colours at the same height to capture bee species (Cane et al. 2000). Although pan trapping does not capture all bee species during the flowering period, it is widely recognized as an efficient method for assessing bee diversity and can detect species that may otherwise be overlooked (Cane et al., 2000). Each trap unit consisted of three coloured bowls/containers (white, blue, and yellow) painted with UV-reflective paint (BOSNY paint). The bowls /containers were filled with water, and approximately 5 mg of unscented washing powder was added to reduce surface tension. Traps were installed for 24 hours. In total, 15 bowls/containers (five clusters) were placed at each of the 19 sampling study sites (a total of 285 bowls/containers) during each sampling bout, giving us 570 bowls/containers across the study period. The average distance between the pan trap stations was a minimum of 1 km.

**Figure 2.**
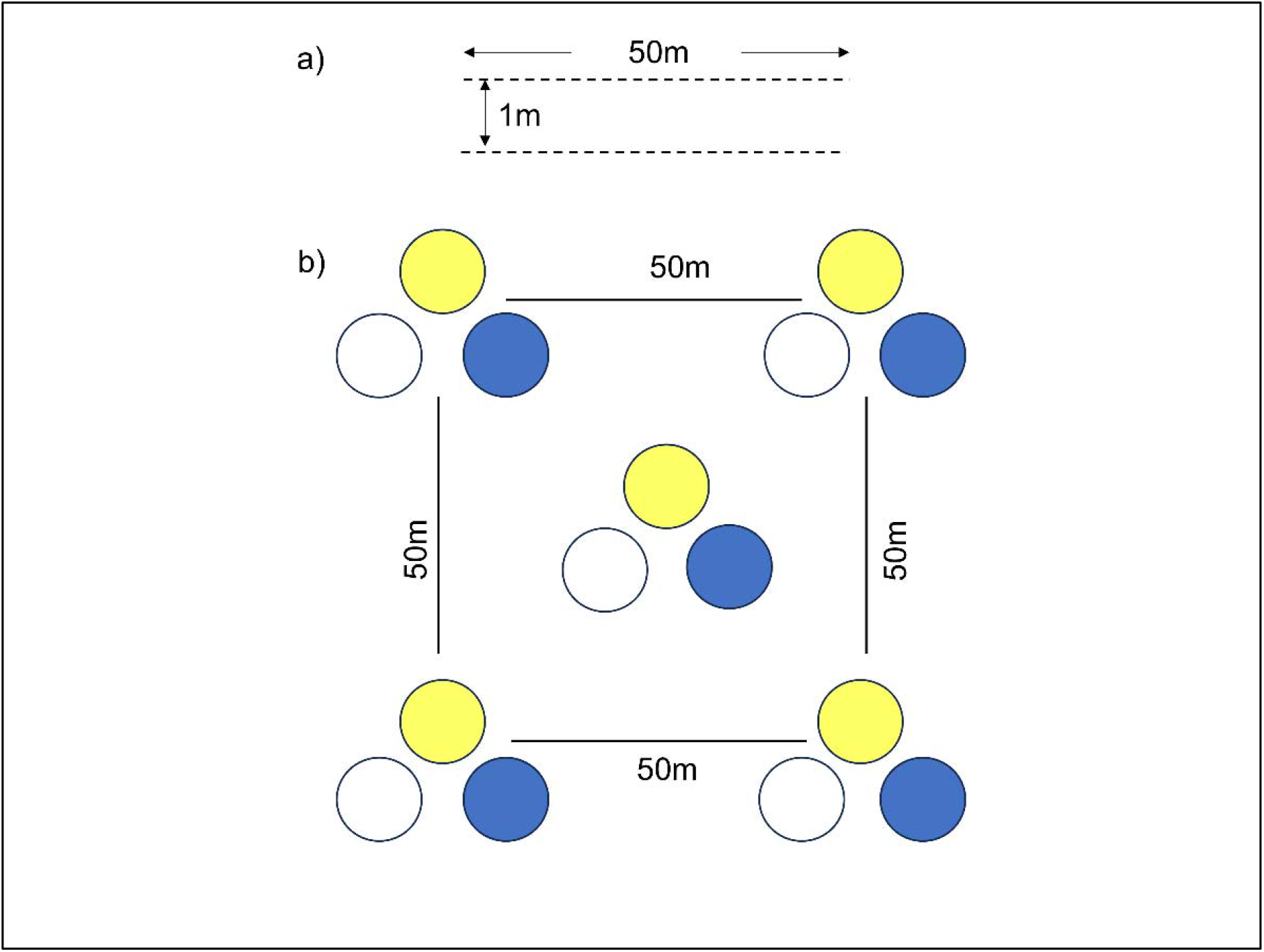
(a) The dotted straight line indicates the transect path in each sampling site, (b) Schematic diagram of the pan trap cluster and colored circles showing the respective colours of pan traps.

For active sampling, bee pollinators were collected using sweep netting. Sampling consisted of 5 minutes of sweeping along a 50 m transect, conducted three times between 8 am and 2 pm by a single collector. The transects were laid parallel to the Pan-trap stations. Collected bees were at first rinsed with distilled water and then preserved in 70% alcohol in the field.

In the laboratory, all the sampled bee specimens were dried, pinned, and identified to the species or morphospecies level using standard taxonomic keys (Michener 2000). Data from the two different sampling rounds and two different sampling methods were pooled for each site for further analysis.

### 2.3. Bee functional trait estimation

For all collected bee species, we gathered information on key life-history and ecological traits. The traits included body size, nesting location, dietary specialization, nest construction behaviour, sociality, hairiness (i.e., density of body hair, which influences pollen deposition on stigmas; Stavert et al., 2017), and glossa length. Glossa length is associated with the range of floral morphologies accessible to bees and influences pollen uptake, transport efficiency, and resource specialization (Michener, 2000; Martins et al., 2015).

Body size was the only quantitative functional trait measured directly. It was estimated using the mean intertegular distance (ITD) of collected specimens, a widely accepted proxy for body size in bees (Greenleaf et al., 2007). For each species, at least ten individuals were measured whenever possible. For species represented by fewer than ten individuals, all available specimens were measured. Information on categorical functional traits was obtained from specialized taxonomic and ecological literature. Species for which trait information could not be reliably assigned were excluded from the functional analyses. A complete list of traits and their classifications is provided in Table 1.

**Table 1.** Traits used in analyses for measuring functional dispersion of bee communities along the high and low AI sites.

| **Trait** | **Categories / Description** | **Source** |
|---|---|---|
| Area of scopa | — | Direct measurement |
| Inter-tegular distance | — | Direct measurement |
| Mouthpart length (mm) | — | Direct measurement |
| Means of pollen transport | (1) Corbiculae; (2) Ventral scopa; (3) Scopa (hind trochanter–basitarsus); (4) Scopa (tibia and basitarsus) | Michener (2007); Viana & Kleinert (2005) |
| Means of pollen uptake | (1) First leg; (2) Vibration; (3) First leg and mandible; (4) First leg and vibration | Michener (2007); Viana & Kleinert (2005) |
| Buzz pollination | Buzz / Non-buzz | Michener (2007); Viana & Kleinert (2005) |
| Peak activity time | (1) Within 0–3 h after sunrise; (2) Within 3–6 h after sunrise | Michener (2007); Viana & Kleinert (2005) |
| Dietary specialization | Oligolectic / Polylectic | Michener (2007); Viana & Kleinert (2005) |
| Sociality | Solitary / Social / Parasitic | Michener (2007); Viana & Kleinert (2005) |
| Nesting location | Above ground / Below ground / Mixed | Michener (2007); Viana & Kleinert (2005) |
| Nest construction | Excavator / Renter / Cleptoparasitic | Michener (2007); Viana & Kleinert (2005) |
| Hairiness (density of hair) | Dense / Sparse | Michener (2007); Viana & Kleinert (2005) |

### 2.4. Calculation of pesticide intensification

Pesticide use data were collected through structured interviews with eight farmers cultivating crops within a 500 m radius of each pan trap station at each study site. For each farm, detailed records were obtained on pesticide type (herbicides, fungicides, and insecticides), commercial name, quantity applied, frequency of application, and treated area during the two months preceding each bee sampling survey. The data collected from the eight farmers at each site were then averaged to obtain site-level estimates of pesticide use. To quantify pesticide pressure, we calculated a Pesticide Intensification (PI) index as the cumulative amount of pesticide applied per unit area (ml m ²) over the two-month period preceding bee sampling, corresponding to the peak cropping phase. This standardized time window was selected to ensure consistent pesticide management data across sites.

### 2.5. Calculation of landscape composition

We recorded the percentage of fallow land (defined as uncultivated fields with very little to no vegetation) and the percentage of surrounding semi-natural vegetation (including forests, hedgerows, woody strips, and grasslands) each study site at 1000m by using Land Use Land Cover (LULC) maps from Sentinel 2 satellite images (ESRI, Inc., Redlands, CA, USA) using the supervised classification method in Arc-GIS software version 10.8.1. Training samples were generated using high-resolution imagery from Google Maps and validated through extensive ground-truthing surveys.

### 2.6. Crop diversity measurement

Crop diversity was recorded within a 500m radius around each 50m x 50m focal site by visiting and observing all 19 study sites during the sampling period with an extensive ground truthing method. The Shannon diversity index for crops was calculated for each study site. Only pollinator dependent crops were surveyed for this study.

### 2.7. Differentiation of study sites by intensification and landscape composition

To separate the 19 sites into high and low agricultural intensification categories, we used Principal Component Analysis (PCA) based on four management and landscape variables: pesticide intensity (ml/m²), crop diversity (Shannon index), percentage semi-natural habitat, and percentage fallow land. Pesticide intensity was considered positively associated with agricultural intensification, whereas crop diversity, semi-natural habitat, and fallow land were considered negatively associated, reflecting more diversified and less intensive farming systems.

All variables were standardized to zero mean and unit variance before analysis, and PCA was performed using the “prcomp” function in R. Agricultural intensification was represented by the scores of the first principal component (PC1), which captured the dominant gradient of variation based on the corresponding variable loadings and their ecological interpretation. PC1 scores were multiplied by –1 so that higher values corresponded to higher agricultural intensification. For comparative analyses, sites were classified into “High” and “Low” agricultural intensification categories using the median of the transformed PC1 scores.

To visualise the distribution of sites in multivariate environmental space, we performed non-metric multidimensional scaling (NMDS) based on Euclidean distances calculated from the same four standardized environmental variables. Differences in environmental composition between the predefined High and Low categories were statistically evaluated using permutational multivariate analysis of variance (PERMANOVA), with 999 permutations. Homogeneity of multivariate dispersion between the categories was assessed using the “betadisper” function.

### 2.8. Statistical analysis

#### 2.8.1. Bee-site network analysis

To investigate differences in bee community composition between high- and low-intensification agricultural sites, we constructed two bee-site networks (bipartite) corresponding to each intensification category. Two separate networks were generated, with one for the high AI sites and the other corresponding to the low AI sites. In the bee-site bipartite framework, sites are treated analogously to plant species in traditional mutualistic networks, where bees utilize site-specific floral and habitat resources, and sites benefit from pollination services provided by bees (Dutta et al., 2024; Basu et al., 2016; Li et al., 2022).

Interaction matrices were developed based on bee abundance across sites within each category. All network analyses were performed using the “bipartite” package in R.

At the network level, structural properties of the bipartite networks were recorded using the “networklevel” function in R. The indices were calculated, including: Connectance, Modularity (Q), Nestedness (NODF), Weighted nestedness (weighted NODF), and Network-level specialization (H2′). Connectance represents the proportion of realized interactions relative to the total number of possible interactions in the network. Modularity (Q) measures the extent to which the network is compartmentalized into distinct interaction modules. Nestedness (NODF) quantifies the degree to which interactions of specialist species form subsets of those of more generalist species. Weighted NODF incorporates interaction frequency into nestedness estimation. The network-level specialization index (H2′) measures the overall degree of interaction specialization across the entire bipartite network and allows comparison between networks with different sizes and interaction densities (Brimacombe et al., 2022).

Species-level metrics were calculated using the “specieslevel” function. For bee species, the following indices were computed: Normalized Degree (ND), Betweenness Centrality (BC), and Species-level specialization (d′). Normalized Degree (ND) quantifies the proportion of realized interactions of a species relative to all possible interactions, indicating its connectivity within the network. Betweenness Centrality (BC) measures how frequently a species lies on the shortest paths connecting other nodes, reflecting its structural importance in maintaining network connectivity. The species-level specialization index (d′), based on Kullback-Leibler distance, quantifies the deviation of observed interactions from random expectations and ranges from 0 (no specialization) to 1 (maximum specialization) (Blüthgen et al. 2006). For site nodes, only the specialization index (d′) was calculated and reported.

#### 2.8.2. Calculation of functional diversity index

To gather information about the functional diversity (FD) of bee morphospecies, we used the function dbFD in package FD, in R version 4.3.3 (2024) with the Cailliez correction for non-Euclidean distances generated by the inclusion of categorical traits (Laliberté et al. 2014) to calculate a few multivariate indices that calculate distinct components of functional diversity, including functional richness (Fric): the volume of multidimensional trait space occupied by the assemblage (Villéger et al., 2008), functional evenness (Feve): the regularity in distribution of the abundance in the functional space (Villéger et al., 2008), functional divergence (Fdiv): the extent to which the pooled abundance is distributed towards the extremes of the functional space of the community (Carmona, 2019), functional dispersion (Fdis): the spread of species and their relative abundance in the functional trait space (Lalibert’e and Legendre, 2010). To calculate functional redundancy (Fred): the degree to which the species in a community occupy the same functional space (Carmona, 2019), we used the uniqueness function in the ‘adiv’ package in R. The value ranged from 0 to 1, with 0 indicating low community redundancy and 1 indicating high community redundancy.

To see the effect of landscape variables and management variables on functional diversity indices, we used GLMs (generalized linear models) with “gaussian” or “Gamma” error distribution and “log” link function based on model fit and distribution of data. To observe any difference in functional diversity indices across high and low AI sites, we performed the Mann-Whitney U test.

#### 2.8.3. Statistical Analysis of Bee Community Metrics

To examine the effects of agricultural management and landscape variables on bee community parameters, we fitted Generalized Linear Models (GLMs) using appropriate error distributions based on the structure of each response variable. Predictor variables were the percentage of semi-natural vegetation, the percentage of fallow land, pesticide intensity (PI), and crop diversity. The relations between total bee abundance and Halictidae family abundance with predictor variables were analysed using a “negative binomial” model with a “log” link function to account for overdispersion in count data. The Shannon bee diversity was analysed using a “gaussian” error distribution with a “log” link function. The other three bee families, Apidae, Andrenidae, and Megachilidae, were excluded from family-level analyses due to their low abundances and absence across most of the study sites.

All models were tested for normality of residuals and homogeneity of variance by Shapiro-wilk test and Levene’s test, respectively, before final selection. The model diagnostics were checked with the “DHARMa” package (Hartig and Hartig, 2017). For all response variables, candidate models were constructed using ecologically relevant predictor variables. Model selection was based on Akaike’s Information Criterion (AIC), and when multiple models were supported (ΔAIC ≤ 2), conditional model averaging was performed to obtain robust parameter estimates.

To observe the difference in bee assemblage between the high and low AI sites, we performed the “Mann-Whitney U” test, taking total bee, bee family abundance, bee diversity, and richness data as the response variables.

## 3. Results

### 3.1. High and low AI gradients

Considerable variation in pesticide intensity and crop diversity was observed across the 19 study sites. Principal Component Analysis (PCA) of the four environmental variables revealed that the first two principal components explained 66.61% of the total variation, with PC1 and PC2 accounting for 37.0% and 29.6%, respectively. PC1 was primarily associated with pesticide intensity (loading = −0.702) and crop diversity (loading = 0.711), indicating that these variables contributed most strongly to the principal axis of variation. Based on the median of the transformed PC1 scores, nine sites were classified as “High” agricultural intensification and ten as “Low” agricultural intensification (Figure 3).

**Figure 3.**
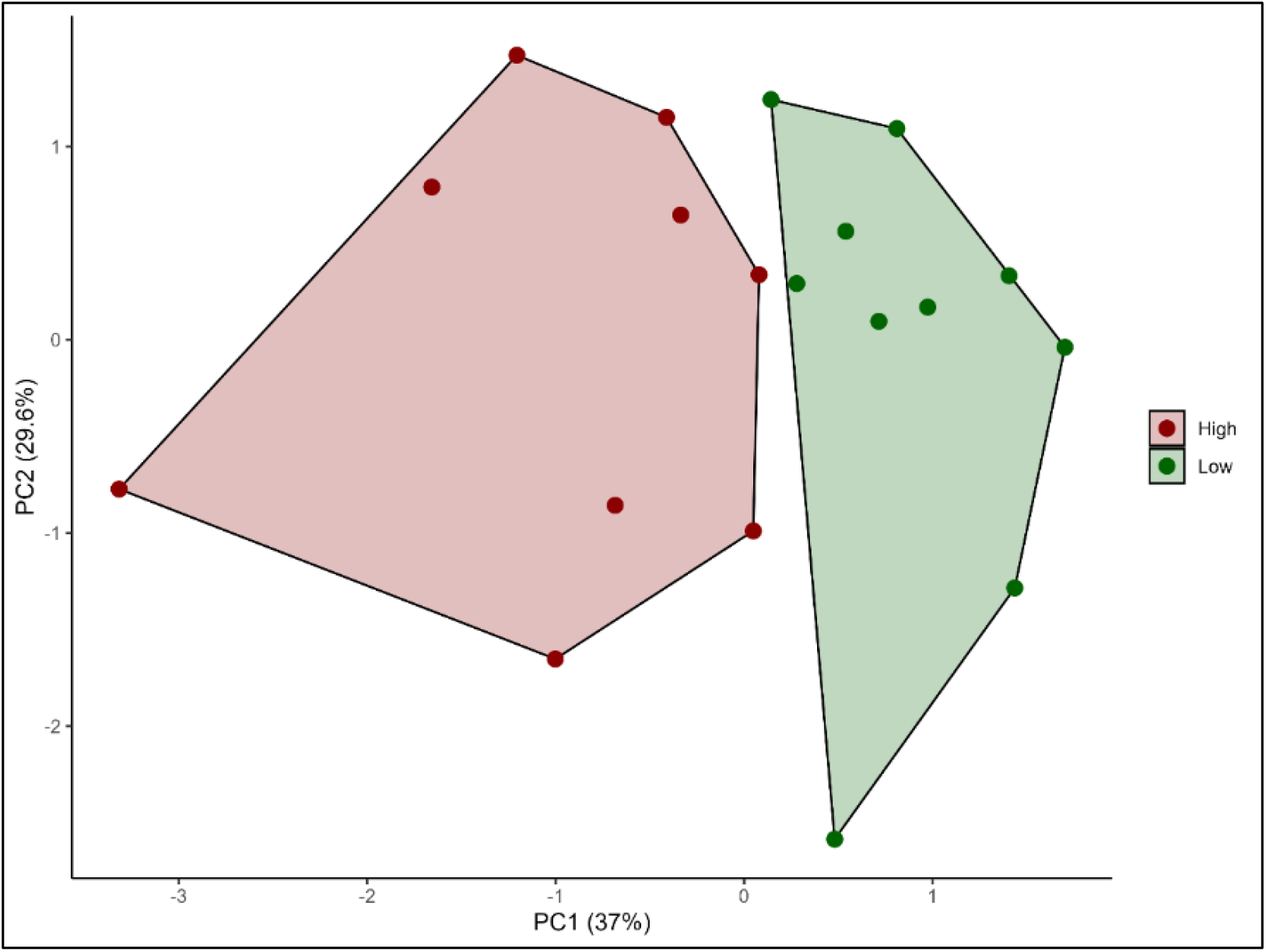
PCA plot showing two separate clusters for High (Red ellipse) and Low (green ellipse) AI sites. Four parameters were taken for the PCA analysis: pesticide index, crop diversity, percentage of semi-natural vegetation, and percentage of fallow-land cover.

Non-metric multidimensional scaling (NMDS), performed using Euclidean distances calculated from the four standardized environmental variables, showed a clear separation of the predefined High and Low agricultural intensification categories (stress = 0.127). All nine High-intensification sites had negative NMDS1 scores, whereas all ten Low-intensification sites had positive NMDS1 scores, indicating complete separation of the two categories along the first ordination axis. PERMANOVA indicated significant differences in multivariate environmental composition between the two categories (F□,□□= 4.99, R² = 0.227, p = 0.001). The homogeneity of multivariate dispersion test revealed no significant difference in within-group dispersion (F□,□□= 0.171, p = 0.663).

Comparisons of individual environmental variables revealed significant differences in crop diversity (W = 2.0, p < 0.001) and pesticide intensity (W = 76.0, p < 0.05) between the two agricultural intensification categories. Pesticide intensity was higher in High-intensification sites (mean ± SD: 0.167 ± 0.08 ml/m²) than in Low-intensification sites (0.100 ± 0.02 ml/m²). Conversely, crop diversity (Shannon index) was lower in High-intensification sites (4.66 ± 1.00) than in Low-intensification sites (6.70 ± 0.94). Although the percentages of semi-natural habitat and fallow land did not differ significantly between the categories, both were higher in Low-intensification sites.

### 3.2. Bee abundance and diversity

A total of 285 bee specimens were found in this study. 11 genera and 22 species were collected across all 19 high and low-intensity study sites during the 2022-2023 sampling sessions. Among the collected bee species, approximately 86% were in the family Halictidae, followed by Andrenidae (6%), Apidae (5%), and Megachilidae (1%). All bee specimens were identified up to the species or morphospecies level. Species richness was higher in the low AI sites (19 species) compared to the high AI sites (10 species). Only one species was unique to the high AI sites, whereas ten species were exclusively recorded in low AI sites. Nine species were shared between the two zones. The unique and common bee species at high and low AI sites are listed in Table 2.

**Table 2.**
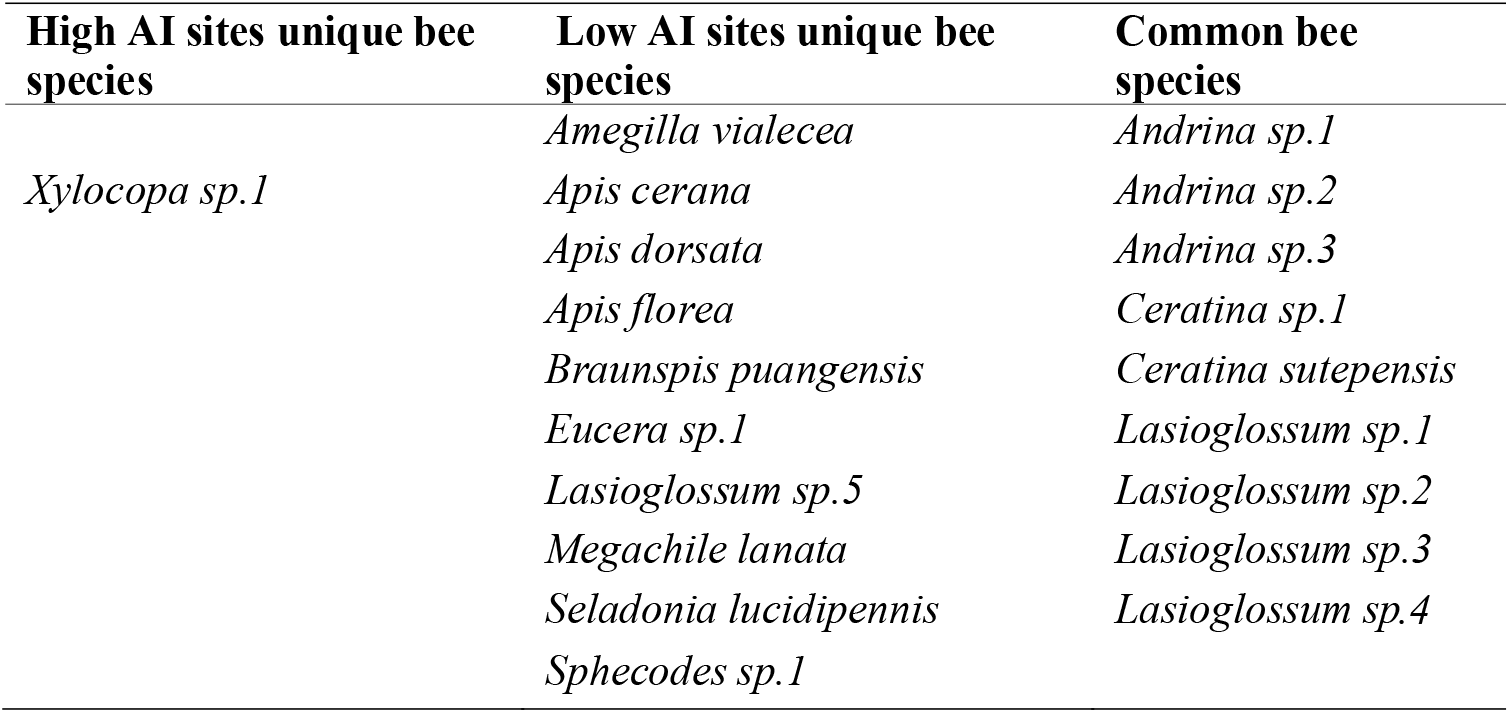
Distribution of bee species across high and low AI sites in the study. High AI areas contained 10 bee species, with 1 species unique to these environments. Low AI areas had 19 species, including 10 unique species. A total of 9 species were common to both areas

| **High AI sites unique bee species** | **Low AI sites unique bee species** | **Common bee species** |
|---|---|---|
|  | *Amegilla vialecea* | *Andrina sp.1* |
| *Xylocopa sp.1* | *Apis cerana* | *Andrina sp.2* |
|  | *Apis dorsata* | *Andrina sp.3* |
|  | *Apis florea* | *Ceratina sp.1* |
|  | *Braunspis puangensis* | *Ceratina sutepensis* |
|  | *Eucera sp.1* | *Lasioglossum sp.1* |
|  | *Lasioglossum sp.5* | *Lasioglossum sp.2* |
|  | *Megachile lanata* | *Lasioglossum sp.3* |
|  | *Seladonia lucidipennis* | *Lasioglossum sp.4* |
|  | *Sphecodes sp.1* |  |

### 3.3. Site bee community network

The bee community networks differed between intensification levels (Figure 4). High AI sites exhibited higher nestedness (NODF = 61) than low AI sites (NODF = 37.49). In contrast, network-level specialization (H′) was slightly higher in low AI sites (H’ = 0.27) compared to high AI sites (H’ = 0.20). All network-level parameters are presented in Table 3.

**Figure 4.**
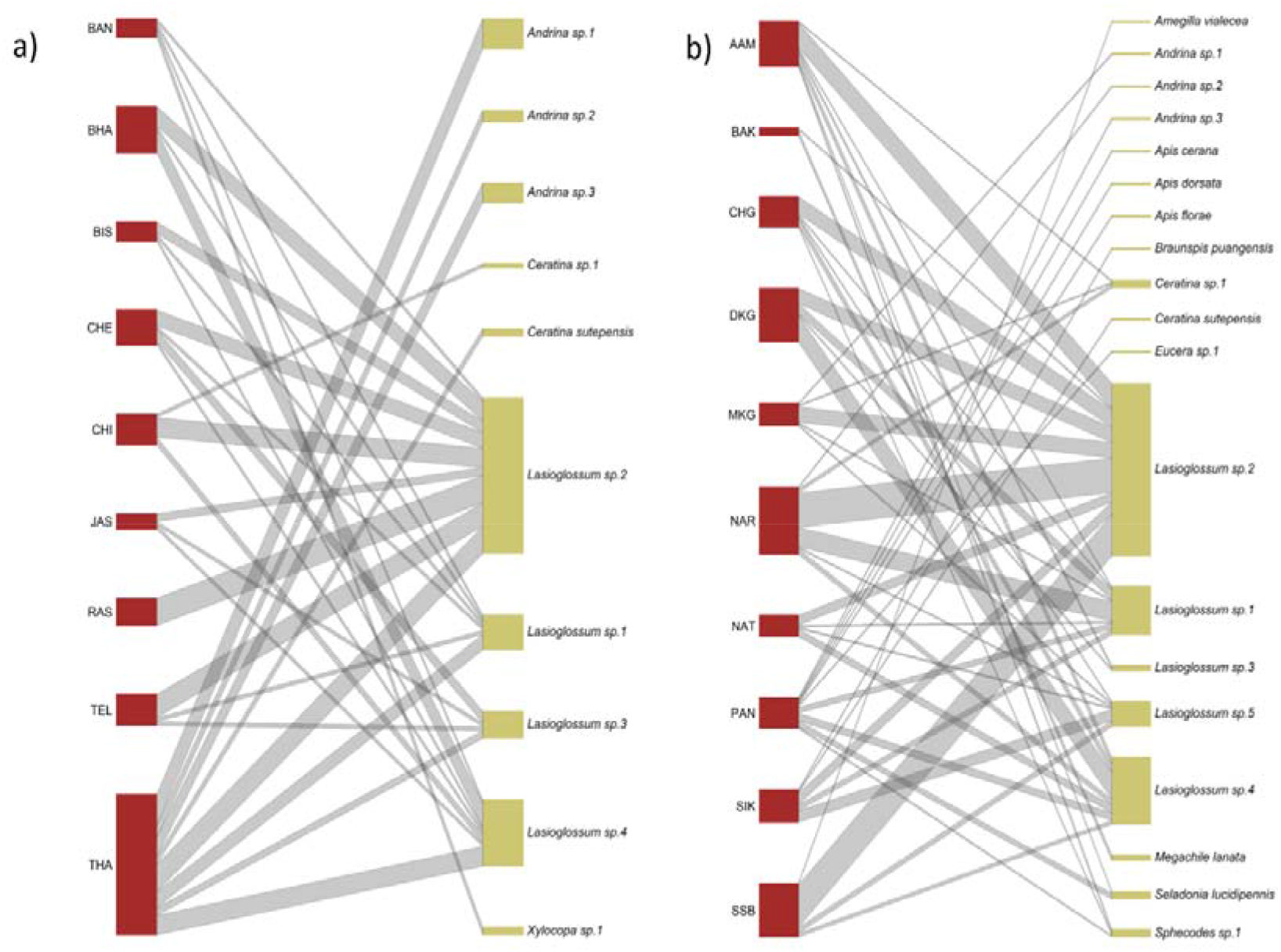
Bee-site networks for high (a) and low (b) clusters. Red boxes represent sites, while yellow boxes denote bee species. The width of each box reflects the relative contribution of each bee species and site to the network. The thickness of the interaction lines corresponds to the number of observed interactions. The bee-site network was developed with R software version 4.3.3

**Table 3.** Bee-site network parameters in low and high AI study sites.

| Metric | Low AI sites | High AI sites |
|---|---|---|
| Connectance | 0.268 | 0.356 |
| Modularity (Q) | 0.276 | 0.321 |
| NODF | 37.50 | 61.01 |
| Weighted NODF | 24.40 | 24.13 |
| H□′ (network-level specialization) | 0.279 | 0.209 |

At the species level, analysis of the site-species matrix showed that *Lasioglossum* sp. 2 had the highest normalized degree (ND) in both high- and low AI sites. The specialization index (d′) was highest for *Megachile lanata* at low AI sites, whereas *Xylocopa* sp. 1 showed the highest d′ value at high AI sites. Betweenness centrality (BC) also varied across regions: at high AI sites, *Lasioglossum* sp. 2 had the highest BC, whereas at low AI sites, *Lasioglossum* sp. 1 had the highest BC.

The mean value of the bee species-specific specialization index did not differ between intensification levels. All bee-level network parameters are provided in Table 4. However, the mean site specialization index was slightly higher at low AI sites (0.23 ± 0.21 SD) than at high AI sites (0.18 ± 0.11 SD) (Table 5).

**Table 4.** Parameters measured at species level for the bee-site network for high and low AI sites. Three parameters at the species level were measured; normalized degree (ND), betweeneness centrality (BC) and species specialization index (d’)

| **Bee species** | **Intensity** | **Normalised degree** | **Betweenness** | **d′** |
|---|---|---|---|---|
| *Andrina sp.1* | high | 0.111111 | 0 | 0.340376 |
| *Andrina sp.2* | high | 0.111111 | 0 | 0.098108 |
| *Andrina sp.3* | high | 0.111111 | 0 | 0.213032 |
| *Ceratina sp.1* | high | 0.111111 | 0 | 0.419721 |
| *Ceratina sutepensis* | high | 0.111111 | 0 | 0.047406 |
| *Lasioglossum sp.2* | high | 1.000000 | 0.430556 | 0.187403 |
| *Lasioglossum sp.1* | high | 0.666667 | 0.138889 | 0.054493 |
| *Lasioglossum sp.3* | high | 0.444444 | 0 | 0.242964 |
| *Lasioglossum sp.4* | high | 0.777778 | 0.430556 | 0.179407 |
| *Xylocopa sp.1* | high | 0.111111 | 0 | 0.698014 |
| *Amegilla vialecea* | low | 0.100000 | 0 | 0.062763 |
| *Andrina sp.1* | low | 0.100000 | 0 | 0.301079 |
| *Andrina sp.2* | low | 0.100000 | 0 | 0.000000 |
| *Andrina sp.3* | low | 0.100000 | 0 | 0.220164 |
| *Apis cerana* | low | 0.100000 | 0 | 0.220164 |
| *Apis dorsata* | low | 0.100000 | 0 | 0.220164 |
| *Apis florae* | low | 0.100000 | 0 | 0.220164 |
| *Braunspis puangensis* | low | 0.100000 | 0 | 0.203112 |
| *Ceratina sp.1* | low | 0.300000 | 0.025167 | 0.150058 |
| *Ceratina sutepensis* | low | 0.100000 | 0 | 0.203112 |
| *Eucera sp.1* | low | 0.100000 | 0 | 0.220164 |
| *Lasioglossum sp.2* | low | 0.900000 | 0.286833 | 0.183198 |
| *Lasioglossum sp.1* | low | 0.800000 | 0.320167 | 0.163473 |
| *Lasioglossum sp.3* | low | 0.200000 | 0 | 0.187650 |
| *Lasioglossum sp.5* | low | 0.600000 | 0.071167 | 0.299939 |
| *Lasioglossum sp.4* | low | 0.700000 | 0.176000 | 0.246789 |
| *Megachile lanata* | low | 0.100000 | 0 | 0.915551 |
| *Seladonia lucidipennis* | low | 0.200000 | 0.050000 | 0.387961 |
| *Sphecodes sp.1* | low | 0.300000 | 0.070667 | 0.215171 |

**Table 5.** Specialization index(d’) measured at the site level for the bee-site network of high and low AI sites. The second column denotes whether the site belongs to a high or low AI cluster.

| **Site** | **Intensity** | **d′** |
|---|---|---|
| BAN | high | 0.334472 |
| BHA | high | 0.201831 |
| BIS | high | 0.022906 |
| CHE | high | 0.155926 |
| CHI | high | 0.203877 |
| JAS | high | 0.022801 |
| RAS | high | 0.274910 |
| TEL | high | 0.157853 |
| THA | high | 0.293314 |
| AAM | low | 0.109015 |
| BAK | low | 0.687243 |
| CHG | low | 0.014885 |
| DKG | low | 0.194893 |
| MKG | low | 0.150532 |
| NAR | low | 0.133303 |
| NAT | low | 0.022887 |
| PAN | low | 0.534339 |

### 3.4. Functional diversity

Agricultural intensification and associated management variables significantly influenced multiple facets of bee functional diversity. Functional richness (FRic) measured from collected bee samples declined significantly with increasing pesticide intensity (p = 0.012) and differed between high- and low AI sites, but the difference was marginally non-significant (*p* = 0.067).

Functional evenness (FEve) of the bee community collected from crop fields decreased with Shannon crop diversity (p = 0.045). Functional dispersion (FDis) significantly decreased with increasing semi-natural vegetation (p = 0.025) and increased with fallow land (marginally non-significant, p = 0.055).

Functional divergence (FDiv) and functional redundancy (FRed) were not significantly associated with the environmental predictors.

### 3.5. Abundance and diversity across high and low AI sites

The Mann-Whitney test showed that total bee abundance (p = 0.036) and Halictidae abundance (p = 0.024) differed significantly between high and low AI sites, with both metrics higher in low AI sites. There was no significant difference between bee diversity across high and low AI sites. We also found no significant difference in the abundance of other bee families between the high and low sites.

### 3.6. Effect of landscape and farm-level management

Total bee abundance and diversity, as well as the responses of major bee families, were significantly associated with landscape and management variables. Total bee abundance decreased significantly with increasing pesticide intensity (p = 0.013). Bee diversity significantly decreased (p = 0.048) with increasing semi-natural vegetation. When bee species were grouped by family, the abundance of Halictidae increased with crop diversity (p = 0.034).

## 4. Discussion

Our study demonstrates that agricultural intensification strongly alters bee community composition, network structure, and functional diversity in a heterogeneous landscape with multiple-crop agricultural farms and semi-natural vegetation. Pesticide intensity emerged as a key driver of bee community assemblage. Total bee abundance declined significantly with increasing pesticide use, indicating that chemical exposure acts as a major ecological stressor. Many studies have found similar results (Bloom et al., 2021; Brittain et al., 2010). Pesticide use at high and low AI sites showed differential effects on bee abundance and the family Halictidae. Several studies have shown that pesticides reduce bee abundance and richness, foraging activity, and nesting behaviour (Raine and Rundlöf, 2024; Nicholson et al., 2024; Schaad et al., 2023; Raine, 2018). Pesticides negatively affect various life-history traits, such as foraging ability, foraging motivation, and colony health (Morandin and Winston 2003; Lämsä et al., 2018). The impact of pesticide exposure varies among bee taxa due to differences in traits such as body size, foraging duration, and degree of specialization. It can further affect population-level parameters, including offspring production and colony growth (Mallinger et al., 2015). Das et al. (2024) further revealed that the loss of pollinators and pollination services due to pesticide usage in crop fields reduced crop production and market gain.

Semi-natural habitats are generally expected to support pollinator communities by providing floral and nesting resources (Kennedy et al., 2013). However, heterogeneous agricultural landscapes do not always guarantee high crop pollination services. For example, studies have shown that even in landscapes with forest cover and diverse habitats, bee abundance can remain low, resulting in inadequate pollination services for crops (Samnegård et al., 2016). This supports our study’s finding that an increasing percentage of semi-natural vegetation is associated with decreased bee diversity in crop fields. In such cases, semi-natural vegetation may serve as refuge habitats where pollinators concentrate their foraging, potentially reducing their visitation to nearby crop fields. Consequently, the presence of semi-natural habitats does not necessarily translate into increased pollinator abundance and diversity within crops, and pollination services may remain limited.

In our study, we found that 86% of bees belong to the family Halictidae, which is significantly higher than in regions with lower crop diversity. We characterized crop diversity by the number of pollinator-dependent crop species. This crop diversity included mass-flowering crops like mustard, which provide good sources of nectar and pollen for bee pollinators and attract large numbers of bees (Holzschuh et al., 2013). Multiple crop types provide more varied and continuous food resources for bees (Schellhorn et al., 2015; Raderschall et al., 2021), which may explain the increase in the abundance of solitary bees from the Halictidae family.

Functional diversity indices of bees collected from crop fields varied with agricultural intensification and landscape composition in our study. Functional richness is the volume of multidimensional trait space occupied by the assemblage. Beyond reductions in abundance, pesticide intensity was associated with a significant decline in functional richness. This pattern indicates that pesticides function as environmental filters, disproportionately affecting functionally distinct or sensitive taxa while allowing disturbance-tolerant generalists to persist. Functional dispersion of bee species collected from crop fields declined with increasing proportions of surrounding semi-natural vegetation. Landscapes containing greater amounts of semi-natural vegetation provide more diverse floral and nesting resources, thereby expanding available niche space (Costa et al., 2017). Under such conditions, many bee species may preferentially forage in these semi-natural habitats, possibly to avoid the effects of agricultural intensification, leaving mainly generalist species within crop fields. Thus, semi-natural vegetation functions as an alternative foraging habitat and a refuge for pollinators (Park et al., 2015). Consistent with this interpretation, Forrest et al. (2015) reported higher bee functional dispersion in natural habitats compared with organic and conventional agricultural systems. Similarly, functional evenness decreased with increasing crop diversity, suggesting that certain functional traits may become dominant in more diverse cropping systems. This pattern could reflect the disproportionate success of a subset of bee species, particularly generalist or disturbance-tolerant taxa, which may exploit available crop resources more efficiently than other functional groups. As a result, trait distribution becomes uneven, even when crop diversity increases. Reduced functional evenness may indicate lower community balance and stability; however, dominant functional groups may still maintain pollination services in agricultural landscapes (Villéger et al., 2008).

Our results revealed clear differences in bee community structure between agricultural intensification levels. In accordance with our hypothesis, high AI sites exhibited higher nestedness compared to low AI sites, indicating that bee communities in highly intensified landscapes were more structured and dominated by generalist species that interact with a broad range of sites. High-intensity management leads to the conversion of complex natural ecosystems into “simplified managed ecosystems”. These intensified landscapes are often “homogeneously farmed” with little non-crop area, which filters out sensitive taxa (REF). These sites, which maintain highly nested networks, are often associated with less heterogeneous habitats in which a few tolerant species persist across multiple locations, whereas more sensitive species occur only in subsets of these communities. This pattern suggests that intensified agricultural landscapes may filter bee communities, favouring disturbance-tolerant and generalist taxa while reducing overall ecological complexity (Tylianakis et al., 2009; Tscharntke et al., 2005). In contrast, lower nestedness at low AI sites indicates a less hierarchical, potentially more heterogeneous community structure.

Network-level specialization (H′) was slightly higher in low AI sites than in high AI sites, suggesting that bee communities in less intensified landscapes exhibited more specialized interactions. This higher specialization may reflect the availability of diverse floral resources, nesting habitats, and microhabitats typically found in low AI systems. Such conditions promote niche differentiation and allow the coexistence of species with varying ecological requirements. Conversely, the lower specialization observed in highly intensified landscapes suggests homogenization of bee communities, with generalist species dominating due to limited resource diversity (Tscharntke et al., 2005).

At the species level, *Lasioglossum* sp. 2 showed the highest normalized degree in both high- and low AI sites, indicating that this species interacted with the greatest number of sites and may function as a generalist species capable of persisting across varying levels of agricultural intensification. Species in the genus *Lasioglossum* are often characterised by ecological flexibility, small body size, and broad floral resource use, which may help explain their dominance across landscapes with varying disturbance levels. Previous studies have similarly reported the dominance of *Lasioglossum* species in similar agricultural landscapes, where soil-nesting bees of this genus increase under agricultural intensification and act as common, disturbance-tolerant taxa (Basu et al., 2016).

The specialization index (d′) further highlighted differences in species roles across intensification gradients. *Megachile lanata* showed the highest specialization in low AI sites, suggesting that this species may depend on specific floral or nesting resources available primarily in less disturbed habitats. In contrast, *Xylocopa* sp. 1 exhibited the highest specialization at high AI sites, indicating that this species may utilize specific resources that remain available even in intensively managed agricultural landscapes, such as woody nesting substrates or particular crop species. Species-level specialisation reflects niche differentiation and resource partitioning, in which species with specific ecological requirements become associated with particular habitats (Dutta et al., 2024). These findings suggest that differences in nesting ecology and resource availability across intensification gradients shape the specialization patterns of bee species.

Betweenness centrality values also varied between intensification levels, indicating differences in species importance for maintaining network connectivity. In high AI sites, *Lasioglossum* sp. 2 exhibited the highest betweenness centrality, suggesting that this species plays a crucial role in connecting different sites and maintaining network stability under intensified conditions. In contrast, *Lasioglossum* sp. 1 showed the highest betweenness centrality in low AI sites, highlighting its importance as a connector species in more heterogeneous landscapes. The dominance of *Lasioglossum* species under intensified conditions may reflect the ecological flexibility of this genus, which enables it to persist across disturbed habitats. Similar patterns have been reported by Basu et al. (2016), where *Lasioglossum* species dominated bee communities and contributed to maintaining network connectivity, with their removal leading to increased specialization and greater network vulnerability. Furthermore, dominance of particular taxa under agricultural intensification is a well-known phenomenon, as increasing intensification and loss of high-quality habitats may shift pollinator communities toward common and ubiquitous species (Carré et al., 2009; Kennedy et al., 2013). These findings suggest that *Lasioglossum* species act as key connector taxa that help maintain network structure and resilience across varying levels of agricultural intensification.

The mean site specialization index was slightly higher in low AI sites, further supporting the idea that less intensified agricultural systems promote more specialized interactions and greater ecological heterogeneity. However, the mean bee species-specific specialization did not differ between intensification levels, suggesting that while site-level specialization varied, individual bee species maintained similar degrees of specialization across landscapes. Overall, these findings highlight that agricultural intensification alters bee community structure, favoring generalist and highly connected species in intensified landscapes while promoting specialization and ecological heterogeneity in low AI systems.

## 5. Conclusion

Our results show that agricultural intensification, especially increased pesticide use, has a strong influence on bee communities in heterogeneous farming landscapes. Sites with higher pesticide input supported fewer bees and lower functional richness, suggesting that these conditions filter out more sensitive species while allowing a smaller set of disturbance-tolerant, generalist taxa to persist. Although semi-natural habitats are often considered beneficial for pollinators, our findings indicate that their presence does not necessarily translate into higher bee diversity within crop fields, possibly because these habitats function as alternative foraging areas.

We also observed clear differences in community structure across intensification levels. Highly intensified sites were characterized by more nested and less specialized interaction patterns, consistent with simplified communities dominated by a few resilient taxa. In contrast, low-intensity systems supported more heterogeneous and specialized interactions, underscoring the role of less disturbed landscapes in maintaining ecological complexity. Changes in functional diversity further suggest that intensification reduces the range of ecological traits present, which could affect the stability of pollination systems over time, even if some level of pollination service is retained.

Overall, our findings revealed that while intensified agricultural systems may retain certain pollination functions, they do so at the cost of reduced biodiversity, functional integrity, and ecological stability. Promoting low-intensity farming practices, reducing pesticide inputs, and managing landscape composition are therefore essential for conserving diverse and resilient bee communities and sustaining pollination services in agricultural landscapes.

## Acknowledgement

We convey our heartfelt regards to the farmers of the study region for allowing us to work in their fields and providing crucial assistance. We also thank Dr. Anirban Chakraborty for helping with the trait measurement. We also thank the Council of Scientific and Industrial Research (CSIR), India, for providing fellowship support to the first author.

## Author Contribution Statement

Dona Roy was involved in conceptualization, fieldwork, formal analysis, writing, review and editing. Indranil Samajpati was involved in conceptualization, formal analysis, and review and editing. Aditi Dutta was involved in formal analysis. Parthiba Basu contributed to the conceptualization and development of the field methodology. Rita Kundu was involved in conceptualization, formal analysis, review and editing.

## Disclosure statements

### Conflicts of interest

The authors declare that there are no conflicts of interest. Research involving human participants and/or animals:

This study did not involve any human or vertebrate animal.

### Informed consent

All surviving authors have reviewed and endorsed the submitted manuscript. The late Parthiba Basu contributed to the study prior to his passing.

## References

Barman, J., Narjinary, S., Biswas, S., Biswas, B. and Jadhav, R., 2022. Elephant habitat suitability analysis of Alipurduar District, West Bengal using geospatial technology. Nature Environment and Pollution Technology, 21(4), pp.1705–1712.

Bascompte, J., Jordano, P., Melián, C.J. and Olesen, J.M., 2003. The nested assembly of plant–animal mutualistic networks. Proceedings of the National Academy of Sciences, 100(16), pp.9383–9387.

Basu, P., Parui, A.K., Chatterjee, S., Dutta, A., Chakraborty, P., Roberts, S. and Smith, B., 2016. Scale dependent drivers of wild bee diversity in tropical heterogeneous agricultural landscapes. Ecology and evolution, 6(19), pp.6983–6992.

Belsky, J. and Joshi, N.K., 2019. Impact of biotic and abiotic stressors on managed and feral bees. Insects, 10(8), p.233.

Bloom, E.H., Wood, T.J., Hung, K.L.J., Ternest, J.J., Ingwell, L.L., Goodell, K.et al. (2021) Synergism between local-and landscape-level pesticides reduces wild bee floral visitation in pollinator-dependent crops. Journal of Applied Ecology, 58(6), 1187–1198. Available from: 10.1111/1365-2664.13871

Blüthgen, N., Menzel, F. and Blüthgen, N., 2006. Measuring specialization in species interaction networks. BMC ecology, 6(1), p.9.

Brimacombe, C., Bodner, K., Michalska Smith, M., Gravel, D. and Fortin, M.J., 2022. No strong evidence that modularity, specialization or nestedness are linked to seasonal climatic variability in bipartite networks. Global Ecology and Biogeography, 31(12), pp.2510–2523.

Brittain, C., and S. G. Potts. 2011. The potential impacts of insecticides on the life-history traits of bees andthe consequences for pollination. Basic Appl. Ecol. 12:321–333.

Brittain, C.A., Vighi, M., Bommarco, R., Settele, J. & Potts, S.G. (2010) Impacts of a pesticide on pollinator species richness at different spatial scales. Basic and Applied Ecology, 11(2), 106–115. Available from: 10.1016/j.baae.2009.11.007

Cane, J.H., Minckley, R.L. and Kervin, L.J., 2000. Sampling bees (Hymenoptera: Apiformes) for pollinator community studies: pitfalls of pan-trapping. Journal of the Kansas entomological society, pp.225–231.

Carmona, C.P., 2019. TPD: Methods for measuring functional diversity based on trait probability density. R package version, 1(0).

Carré, G., Roche, P., Chifflet, R., Morison, N., Bommarco, R., Harrison-Cripps, J., Krewenka, K., Ponisio, L., Potts, S.G., Roberts, S.P.M., Schweiger, O., Smith, H.G., Steffan-Dewenter, I., Szentgyörgyi, H., Tscheulin, T. & Westphal, C., 2009. Landscape context and habitat type as drivers of bee diversity in European annual crops. Agriculture, Ecosystems & Environment, 133(1–2), pp.40– 47.

Chakraborty, A. and Basu, P., 2025. Intensive agriculture influences functional diversity, redundancy and trait profile of bee community and interacting plant community in a tropical agricultural landscape. Agriculture, Ecosystems & Environment, 383, p.109544.

Coutinho, J.G., Hipólito, J., Santos, R.L., Moreira, E.F., Boscolo, D. and Viana, B.F., 2021. Landscape structure is a major driver of bee functional diversity in crops. Frontiers in Ecology and Evolution, 9, p.624835.

Das, M., Paul, A., Bhattacharya, S. and Basu, P., 2025. Pollinator visitation decline due to pesticide application beyond threshold frequency brings down crop yield. Ecological Entomology, 50(1), pp.33–42.

Das, T., 2020. Socio-econmic characteristics of forest villagers in Alipurduar district of West Bengal (Doctoral dissertation, University of North Bengal).

Dong, Z., Bladon, A.J., Jaworski, C.C., Pywell, R.F., Woodcock, B.A., Meek, W.R., Nuttall, P. and Dicks, L.V., 2025. Species habitat networks reveal conservation implications that other community analyses do not detect. Ecological Applications, 35(1), p.e3073.

Dutta, A., Samajpati, I. and Basu, P., 2024. Bee community response to multiple stressors along a tropical urban-peri urban gradient. Urban Ecosystems, 27(6), pp.2569–2586.

Fahrig, L. 2003. Effects of habitat fragmentation on biodiversity. Annu. Rev. Ecol. Evol. Syst. 34:487–515.

Felderhoff, J., Gathof, A.K., Buchholz, S. and Egerer, M., 2023. Vegetation complexity and nesting resource availability predict bee diversity and functional traits in community gardens. Ecological Applications, 33(2), p.e2759.

Fletcher Jr, R.J., Revell, A., Reichert, B.E., Kitchens, W.M., Dixon, J.D. and Austin, J.D., 2013. Network modularity reveals critical scales for connectivity in ecology and evolution. Nature communications, 4(1), p.2572.

Forrest, J.R., Thorp, R.W., Kremen, C. and Williams, N.M., 2015. Contrasting patterns in species and functional trait diversity of bees in an agricultural landscape. Journal of Applied Ecology, 52(3), pp.706–715.

Gámez-Virués, S., Perović, D.J., Gossner, M.M., Börschig, C., Blüthgen, N., De Jong, H., Simons, N.K., Klein, A.M., Krauss, J., Maier, G. and Scherber, C., 2015. Landscape simplification filters species traits and drives biotic homogenization. Nature communications, 6(1), p.8568.

Garibaldi, L.A., Steffan Dewenter, I., Kremen, C., Morales, J.M., Bommarco, R., Cunningham, S.A., Carvalheiro, L.G., Chacoff, N.P., Dudenhöffer, J.H., Greenleaf, S.S. and Holzschuh, A., 2011. Stability of pollination services decreases with isolation from natural areas despite honey bee visits. Ecology letters, 14(10), pp.1062–1072.

Geiger, F., J. Bengtsson, F. Berendse, W. W. Weisser, M. Emmerson, M. B. Morales, et al. 2010. Persistent negative effects of pesticides on biodiversity and biological control potential on European farmland. Basic Appl. Ecol. 11:97– 105.

Ghosh, S., Mandal, M., Vijayaprakash, V., Pandey, D. and Gayen, S.K., 2025. Geospatial analysis of elephant mortalities by electrocution from northern districts landscape of West Bengal, India. Environmental Monitoring and Assessment, 197(10), p.1124.

Greenleaf, S.S., Williams, N.M., Winfree, R. and Kremen, C., 2007. Bee foraging ranges and their relationship to body size. Oecologia, 153(3), pp.589–596.

Hartig, F. and Hartig, M.F., 2017. Package ‘dharma’. R package, 531, p.532.

Holzschuh, A., Dormann, C.F., Tscharntke, T. and Steffan-Dewenter, I., 2013. Mass-flowering crops enhance wild bee abundance. Oecologia, 172(2), pp.477–484.

Kennedy, C. M., E. Lonsdorf, M. C. Neel, N. M. Williams, T. H. Ricketts, R. Winfree, et al. 2013. A global quantitative synthesis of local and landscape effects on wild bee pollinators in agro ecosystems. Ecol. Lett. 16:584–599.

Kennedy, C.M., Lonsdorf, E., Neel, M.C., Williams, N.M., Ricketts, T.H., Winfree, R., Bommarco, R., Brittain, C., Burley, A.L., Cariveau, D. and Carvalheiro, L.G., 2013. A global quantitative synthesis of local and landscape effects on wild bee pollinators in agroecosystems. Ecology letters, 16(5), pp.584–599.

Klein, A. M., B. E. Vaissiere, J. H. Cane, I. Steffan-Dewenter, S. A. Cunningham, C. Kremen, et al. 2007. Importance of pollinators in changing landscapes for world crops. Proc. Biol. Sci. 274:303–313.

Klein, A.M., Vaissière, B.E., Cane, J.H., Steffan-Dewenter, I., Cunningham, S.A., Kremen, C. and Tscharntke, T., 2007. Importance of pollinators in changing landscapes for world crops. Proceedings of the royal society B: biological sciences, 274(1608), pp.303–313.

Laliberté, E. and Legendre, P., 2010. A distance based framework for measuring functional diversity from multiple traits. Ecology, 91(1), pp.299–305.

Laliberté, E., Legendre, P., Shipley, B. and Laliberté, M.E., 2014. Package ‘fd’. Measuring functional diversity from multiple traits, and other tools for functional ecology, 1, pp.0–12.

Lami, F., I. Bartomeus, D. Nardi, T. Beduschi, F. Boscutti, P. Pantini, G. Santoiemma, C. Scherber, T. Tscharntke, and L. Marini. 2021. “Species-Habitat Networks Elucidate Landscape Effects on Habitat Specialisation of Natural Enemies and Pollinators.” Ecology Letters 24: 288–297.

Lasway, J.V., Peters, M.K., Njovu, H.K., Eardley, C., Pauly, A. and Steffan Dewenter, I., 2022. Agricultural intensification with seasonal fallow land promotes high bee diversity in Afrotropical drylands. Journal of Applied Ecology, 59(12), pp.3014–3026.

Li, N., Yang, X., Ren, Y. and Wang, Z., 2022. Importance of species traits on individual-based seed dispersal networks and dispersal distance for endangered trees in a fragmented forest. Frontiers in plant science, 13, p.1010352.

Marini, L., Bartomeus, I., Rader, R. and Lami, F., 2019. Species–habitat networks: A tool to improve landscape management for conservation. Journal of Applied Ecology, 56(4), pp.923–928.

Marini, L., I. Bartomeus, R. Rader, and F. Lami. 2019. “Species Habitat Networks: A Tool to Improve Landscape Management for Conservation.” Journal of Applied Ecology 56: 923–28.

Martins, K.T., Gonzalez, A. and Lechowicz, M.J., 2015. Pollination services are mediated by bee functional diversity and landscape context. Agriculture, Ecosystems & Environment, 200, pp.12–20.

Michener CD (2000) The bees of the world, vol 1. JHU

Michener, C.D., 2007. The Bees of the World Johns Hopkins University Press. Baltimore, Md, USA.

Nicholls, C.I. and Altieri, M.A., 2013. Plant biodiversity enhances bees and other insect pollinators in agroecosystems. A review. Agronomy for Sustainable development, 33(2), pp.257–274.

Park, M.G., Blitzer, E.J., Gibbs, J., Losey, J.E. and Danforth, B.N., 2015. Negative effects of pesticides on wild bee communities can be buffered by landscape context. Proceedings of the Royal Society B: Biological Sciences, 282(1809).

Potts, S.G., Biesmeijer, J.C., Kremen, C., Neumann, P., Schweiger, O. and Kunin, W.E., 2010. Global pollinator declines: trends, impacts and drivers. Trends in ecology & evolution, 25(6), pp.345–353.

Potts. 2010. Impacts of a pesticide on pollinator species richness at different spatial scales. Basic Appl. Ecol. 11:106– 115.

Priyadarshana, T.S., Martin, E.A., Sirami, C., Woodcock, B.A., Goodale, E., Martínez Núñez, C., Lee, M.B., Pagani Núñez, E., Raderschall, C.A., Brotons, L. and Rege, A., 2024. Crop and landscape heterogeneity increase biodiversity in agricultural landscapes: A global review and meta analysis. Ecology Letters, 27(3), p.e14412.

Raderschall, C.A., Bommarco, R., Lindström, S.A. and Lundin, O., 2021. Landscape crop diversity and semi-natural habitat affect crop pollinators, pollination benefit and yield. Agriculture, Ecosystems & Environment, 306, p.107189.

Regmi, B., Dahal, S., Poudel, S.R. and Thomas, P.T., 2025. Native pollinators in Nepal: their importance, threats, and conservation needs. Global Ecology and Conservation, p.e03934.

Robinson, R. A., and W. J. Sutherland. 2002. Post-war changes in arable farming and biodiversity in Great Britain. J. Appl. Ecol. 39:139–176.

Samnegård, U., Hambäck, P.A., Lemessa, D., Nemomissa, S. and Hylander, K., 2016. A heterogeneous landscape does not guarantee high crop pollination. Proceedings of the Royal Society B: Biological Sciences, 283(1838).

Schellenberger Costa, D., Gerschlauer, F., Pabst, H., Kühnel, A., Huwe, B., Kiese, R., Kuzyakov, Y. and Kleyer, M., 2017. Community weighted means and functional dispersion of plant functional traits along environmental gradients on Mount Kilimanjaro. Journal of Vegetation Science, 28(4), pp.684–695.

Schellhorn, N.A., Gagic, V. and Bommarco, R., 2015. Time will tell: resource continuity bolsters ecosystem services. Trends in ecology & evolution, 30(9), pp.524–530.

Singh, N.P., Anand, B., Singh, S., Srivastava, S.K., Rao, C.S., Rao, K.V. and Bal, S.K., 2021. Synergies and trade-offs for climate-resilient agriculture in India: an agro-climatic zone assessment. Climatic change, 164(1), p.11.

Stavert, J.R., Pattemore, D.E., Gaskett, A.C., Beggs, J.R. and Bartomeus, I., 2017. Exotic species enhance response diversity to land-use change but modify functional composition. Proceedings of the Royal Society B: Biological Sciences, 284(1860).

Steffan-Dewenter, I., and T. Tscharntke. 2000. Butterfly community structure in fragmented habitats. Ecol. Lett. 3:449–456.

Stouffer, D.B. and Bascompte, J., 2011. Compartmentalization increases food-web persistence. Proceedings of the National Academy of Sciences, 108(9), pp.3648–3652.

Tscharntke, T., Clough, Y., Wanger, T.C., Jackson, L., Motzke, I., Perfecto, I., Vandermeer, J. and Whitbread, A., 2012. Global food security, biodiversity conservation and the future of agricultural intensification. Biological conservation, 151(1), pp.53–59.

Tscharntke, T., Klein, A.M., Kruess, A., Steffan Dewenter, I. and Thies, C., 2005. Landscape perspectives on agricultural intensification and biodiversity–ecosystem service management. Ecology letters, 8(8), pp.857–874.

Tylianakis, J.M., Laliberté, E., Nielsen, A. and Bascompte, J., 2010. Conservation of species interaction networks. Biological conservation, 143(10), pp.2270–2279.

Viana, B.F. and Kleinert, A.D.M.P., 2005. A community of flower-visiting bees (Hymenoptera: Apoidea) in the coastal sand dunes of northeastern Brazil. Biota neotropica, 5, pp.79–91.

Villéger, S., Mason, N.W. and Mouillot, D., 2008. New multidimensional functional diversity indices for a multifaceted framework in functional ecology. Ecology, 89(8), pp.2290–2301.

